# Software for Non-Parametric Image Registration of 2-Photon Imaging Data

**DOI:** 10.1101/2021.07.25.453381

**Authors:** Philipp Flotho, Shinobu Nomura, Bernd Kuhn, Daniel J. Strauss

**Affiliations:** Systems Neuroscience and Neurotechnology Unit, Neurocenter, Faculty of Medicine, Saarland University & School of Engineering, htw saar, Germany; Summer Program, Japan Society for the Promotion of Science (JSPS), Tokyo, Japan; Optical Neuroimaging Unit, Okinawa Institute of Science and Technology Graduate University, Tancha, Onna-son, Kunigami, Okinawa, Japan

## Abstract

Functional 2-photon microscopy is a key technology for imaging neuronal activity. The recorded image sequences, however, can contain non-rigid movement artifacts which requires high-accuracy movement correction. Variational *optical flow* (OF) estimation is a group of methods for motion analysis with established performance in many computer vision areas. However, it has yet to be adapted to the statistics of 2-photon neuroimaging data. In this work, we present the motion compensation method *Flow-Registration* that outperforms previous alignment tools and allows to align and reconstruct even low *signal-to-noise ratio* 2-photon imaging data and is able to compensate high-divergence displacements during local drug injections. The method is based on statistics of such data and integrates previous advances in variational OF estimation. Our method is available as an easy-to-use ImageJ / FIJI plugin as well as a MATLAB toolbox with modular, object oriented file IO, native multi-channel support and compatibility with existing 2-photon imaging suites.

## 1 Introduction

2-photon microscopy in combination with synthetic or genetically encoded indicators allows to image a wide range of different aspects of neuronal activity with cellular or even sub-cellular resolution in anesthetized as well as behaving animals [1, 2]. Importantly, small signal changes might carry crucial information. However, the imaging data can be afflicted with different types of noise and artifacts. On one hand, due to the low number of generated photons with 2-photon excitation, the shot noise is typically significant. On the other hand, movement noise can be introduced during acquisition. Motion artifacts can be caused by heart beat, breathing, as well as motor behavior in awake animals. Also, some experimental paradigms inherently result in large, non-rigid and non-elastic deformations, for example, local drug injections. While many established tools exist for the correction of small vibrations and rigid drifts, the compensation of large and/or non-uniform motion is still a challenge.

Furthermore, for high accuracy alignment, subpixel registration is necessary: Small displacements can be approximated by a linearized motion assumption [3]. This means even small residual motion can induce large artifacts around image edges due to a proportional relation of spatial image gradient and motion magnitude with respect to induced brightness changes.

While *optical flow* (OF) based image registration methods were used for 2-photon imaging data before [4], they do not incorporate the advances in OF estimation developed in recent years in the context of many computer vision areas.

As a consequence, they perform poorly, when compared with state-of-the-art image registration methods for 2-photon imaging and are generally regarded as too prone to noise for this application [5].

There has been a great deal of work on OF techniques in the past decades with the goal of improving accuracy, model invariants as well as robustness and computing speed. The recent advancements in OF estimation tackle the problem of large displacements with discontinuous motion fields and motion layers. Chen et al. [6] lead (as of July 2021) the Middlebury optical flow benchmark [7] with respect to *average endpoint error* (AEE) and *average angular error* (AAE). They use similarity transformations for a segmented flow field as initialization of large motion. In a second step, the variational method of Sun et al. [8] is used for subpixel refinement.

In this work, we propose a novel, OF based image registration approach for the motion compensation of 2-Photon neuroimaging data. We build on the well studied framework for variational OF estimation [3, 9, 8] and adapt this framework to 2-photon imaging data by techniques which have recently been developed in visual computing. We demonstrate the performance on challenging 2-photon imaging data and can report state-of-the-art results in terms of registration quality, competing computation speed and easy accessibility. Our method is available as an easy-to-use MATLAB toolbox as well as an ImageJ / FIJI plugin.

## 2 Materials and Methods

In this section, we will first briefly describe the acquisition methods for the data that is used in the benchmark dataset and then present the theory as well as implementation details of the proposed motion compensation method Flow-Registration.

### 2.1 Animals

Experiments were approved by the OIST Institutional Animal Care and Use Committee, and performed in and AAALAC International accredited facility. Animals were maintained in a 12 h / 12h light/dark cycle at 22 °C, with food and water available *ad libitum*.

### 2.2 Recombinant viruses

The adeno-associated virus (AAV) encoding the GAkdYmut PKA activity sensor [10] was custom made and produced by the vector core facility of Pennsylvania University (AAV2/1-hSyn-GAkdYmut-hGH, titer: 4 × 1014 gc/ml), and mixed with AAV2/1- hSyn-TurboRFP-WPRE encoding the red fluorophore RFP (titer: 4 × 1013 gc/ml, same supplier) at a ratio 1 : 1.

### 2.3 Expression of GAkdYmut and TurboRFP in cortex

Viral transfer of the indicator gene into cortical neurons of the mouse was performed as described before (Nomura et al. [11]). C57/BL6 mice (2-month-old) were anesthetized with a mixture of medetomidine (0.3mg/kg), midazolam (4mg/kg) and butorphanol (5mg/kg). After performing a craniotomy over somatosensory cortex (AP −1.5 mm, ML 1.7mm, DV –0.6-0.7mm from bregma), 70-140 nL of a 1 : 1 mixture of AAV2/1- hSyn-GAkdYmut-hGH and AAV2/1-hSyn-TurboRFP-WPRE was injected in layer V at a rate of 10 nL/min. A chronic cranial window with a silicon access port (5 mm glass coverslip) was mounted as described Roome and Kuhn [12, 13]. At the end of the surgery, mice received atipamezole (0.3mg/kg) for recovery from anesthesia, and buprenorphine (0.1 mg/kg) for pain relieve. Five to eight weeks after the AAV injection, mice were head-fixed for imaging experiments performed under anesthesia with 1% isoflurane or awake.

### 2.4 In vivo imaging in cortex

A combined wide-field / two-photon microscope (MOM, Sutter Instruments) with a femtosecond-pulsed Ti:sapphire laser (Vision II, Coherent) was used. To increase the point spread function of excitation the back aperture of the 25 × water immersion objective (Olympus) was underfilled (spatial resolution 1μm × 1μm × 4ιm). The collar of the objective was adjusted to correct for the window glass thickness (170 μm). Simultaneous excitation of GAkdYmut (GFP-based single fluorophore sensor, Bonnot et al. [10]) and TurboRFP was performed at 950 nm with a typical power of 5-11 mW. Fluorescence was detected in two channels by GaAsP photomultipliers (Hamamatsu) in spectral windows 490–550 nm (GAkdYmut) and 600-700 nm (TurboRFP), separated by a 560 nm dichroic mirror (all Semrock). The microscope was controlled by a commercial software (MScan, Sutter Instruments). Sampling rate was 30.9 frame/s with 512 × 512 pixel, corresponding to a field of view of 375μm × 375 μm. Saline (0.9% NaCl) with Alexa592 (1μM) or with additional drug (propranolol 10 mM) was injected under anesthesia. Pressure injection was performed through the silicon access port using a glass pipette beveled to a diameter of 5-10 μm opening.

### 2.5 Motion compensation

The motion statistics for 2-photon imaging differ from the motion we encounter in natural images in several ways: Due to the illumination strategy, there is only a single imaging plane and thus no different motion layers. Also, the imaged object is usually soft, biological tissue and, as a consequence, we expect smooth motion fields without discontinuities. We expect the image to contain a fixed *region of interest* (ROI) with small displacements between frames and, potentially, large drift over time. Due to the nature of the scanning method, horizontal displacements may occur. Also, the images are usually not represented in perceptual color spaces, restricting the use of classical photometric invariances, such as in [14], but often there exist multiple signal and or structural channels. On top of that, due to technical limitations, high speed imaging can often only be realized on narrow field of views (FOV).

Our image registration method builds on those observations: Due to the absence of different motion layers as well as of large displacements, we do neither need strategies for large displacements as in [6] nor a regularizer that preserves discontinuities as in [8]. Also, the assumptions of elastic regularizers which penalize the divergence of the OF field, e.g. compare [15], are violated by recordings during drug injection, see Figure 1 (F), where the displacement field has a high divergence after injection. Therefore, we quadratically penalize deviations from smooth displacements fields. In our previous work [16], we developed a motion compensation strategy for 1D linescans (e.g. [17]). We extend this approach to compensate recordings with narrow FOV and implement non-uniform warping where we perform more warping steps along the larger image dimension during optimization.

**Figure 1:**
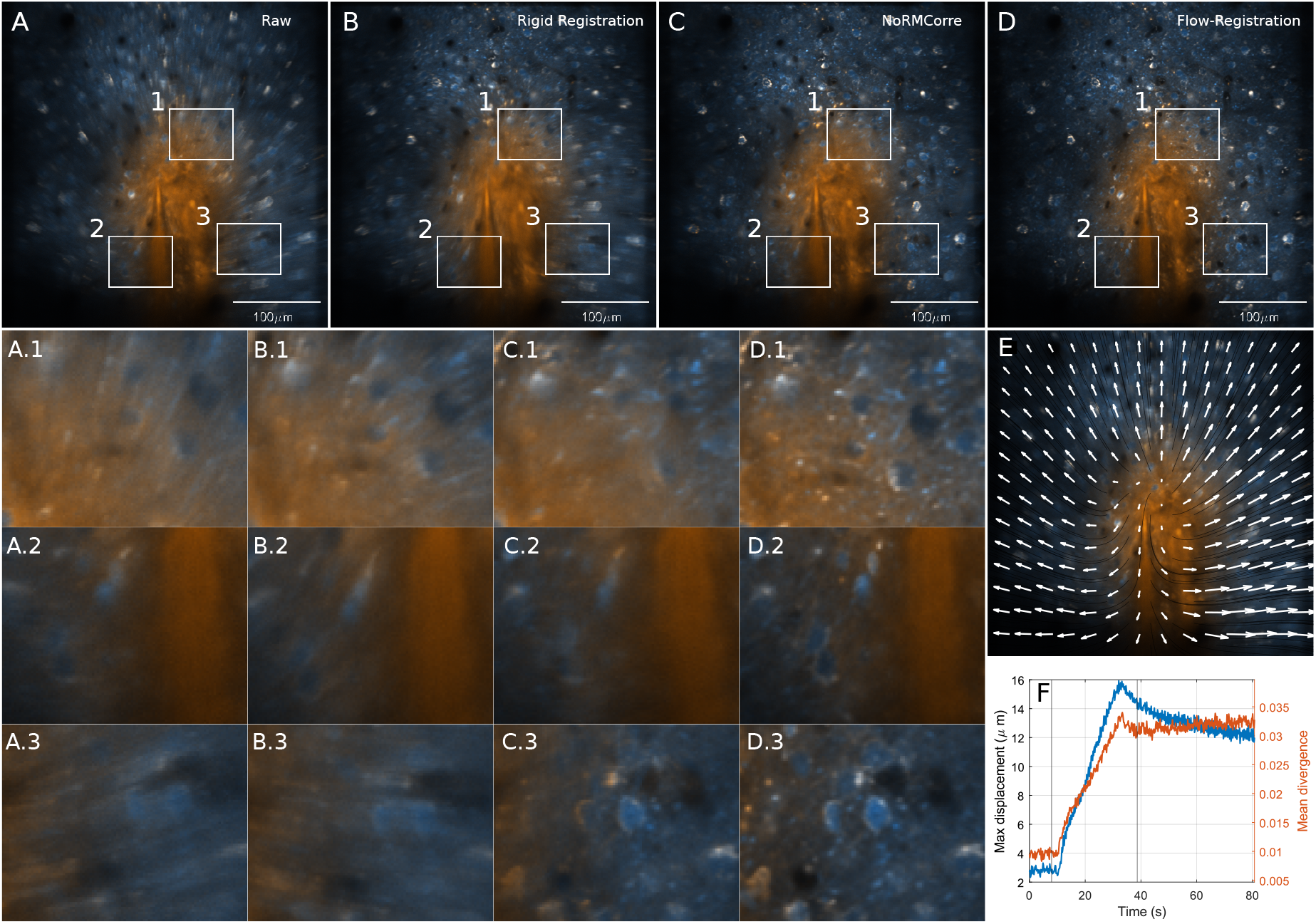
Comparison of registration performance on a very challenging two-channel 2-photon recording during drug injection in vivo. The challenges of the sequence are a very low signal-to-noise ratio together with brightness changes in the functional imaging channel (orange) and non-elastic deformations due to the injected indicator. Average of (A) raw images, (B) after rigid registration, (C) after registration with NoRMCorre, and (D) after Flow-Registration. The tissue expands from the injection point (E) resulting in large displacements as well as in a high divergence (F) in the displacement field. Flow-Registration can recover fine structures with much more detail, allowing region-of-interest selection along fine structures, while the blur and double images in (A-C) indicate residual motion. The channels in the false color representation have all been normalized with respect to the min and max intensity values of the averaged raw recording.

We model this as a robust variational OF method with gradient constancy, robust, separate channel penalization of the dataterm and first-order, isotropic, flow-driven regularizer. We normalize the dataterm according to Zimmer et al. [14] and apply the recommended good practices proposed by Sun et al. [8]. The regularizer and dataterm are penalized with a generalized Charbonnier penalty function which for *ε* > 0 is given by Ψ_*a*_(*x*) = (*x*^2^ + *ϵ*^2^)^*a*^, for *a* > 0 [8]. For the regularizer, larger values of *a* reduce discontinuities in the flow field (staircasing artifacts) and for the dataterm, smaller values of a reduce the influence of outliers. For 2-Photon recordings, we set the smoothness a to 1, which results in quadratic penalization. Note, that for *a* = 0.5 we get a regularized ℓ^1^ norm, while the function becomes non-convex for values of *a* < 0.5. However, Sun et al. encourage a choice of *a* = 0.45 on the Middlebury benchmark, which we apply for the dataterm here. For the numerical approximation, we follow the framework of Bruhn et al. [18], Brox et al. [9] and Papenberg et al. [19] and discretize the *Euler Lagrange equations* in the compact *motion tensor* notation to solve them with an iterative multiscale solver (downsampling factor *η* ∈ (0,1)) with lagged non-linearities with an update in every 5 iterations. We use bicubic interpolation for the warping steps and 5 × 5 median-filtering (mirror boundary) of the flow increments for each level to increase accuracy as suggested by Sun et al. [8].

Low-pass filtering is an important pre-processing step for local and global methods [20]. It makes the images derivable and integrates over temporal changes, while removing image noise. We found convolution with a 3D Gaussian kernel together with subquadratic penalization, to be robust enough to deal with the noise in the benchmark data, e.g., compare the layer 5 data (see supplemental Figure 2). Previous work has shown that registration on structural channels does not necessarily result in much better registration performance [5]. However, in the variational framework, we minimize the joint energy of the structural and the functional channels.

This has the advantage that, in theory, we get a better SNR, if the same structures are visible in both channels and, additionally, considering the aperture proplem, we have potentially more information on the motion whenever disjunct structures are visible in the different image channels, such as in the data in supplemental Figure 2 (A)-(D). Additionally, the energy functional can naturally incorporate a manual weighting term per channel, such as ROIs, to enforce a low value of the dataterm around important image structures. In our implementation, a spatial weight term can be supplied for this purpose.

The main parameters of our method are the regularization parameter *α* = (*α*_1_, *α*_2_)^⊤^, the penalizer power *a*, the warping depth, the downsampling factor *η* and for each channel the Gaussian kernel size *σ* = (*σ*_1_, *σ*_2_, *σ*_3_) for pre-processing. The choice of *α* presents the compromise of correctly registering smaller structures that deviate from the global motion direction and a globally smooth solution. With different values for *α*_1_ and *α*_2_, the smoothness term becomes anisotropic. This might improve the estimation of motion artifacts induced by horizontal scanning. We scale alpha on each level of the solver scale space such that at a level *i* we use *α* = *α* · *η* ^−*i*/2^. In practice, this made the result more robust under lower choices of α while enabling the compensation of high frequent jitter.

For OI, the recorded image section is usually fixed. Therefore, we use a fixed reference frame *f*_ref_ and estimate the flow field *w_t_* = (*u_t_, v_t_*)^⊤^ at time *t* such that *f*_ref_(*x* + *u_t_, y* + *v_t_*) depicts the same scene points as *f*(*x,y,t*). For the final motion compensation step, we apply backward warping with bilinear interpolation to calculate the motion compensated frame at time *t* as 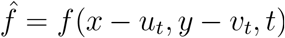.

### 2.6 Toolbox design and implementation

The ImageJ / FIJI plugin makes use of the ImageJ file formats and therefore can resort to all supported file types. Our software design for the MATLAB toolbox consists of modular file readers and writer classes that can be automatically instantiated or supplied as parameters to an options object that defines registration jobs. By default, MDF, tiff image stacks, MATLAB mat files or hierarchical dataformat (HDF5) files are supported. Additionally, the output formats support HDF5 files that are readable by the 2-Photon imaging suites Begonia[21] and CaImAn[22]. The file IO is designed for multichannel processing and supplies 4D matrices in the format height × width × channel × time independently of the actual data representation on disk. To compensate a recording, the file reader supplies batches of size *n* with on-the-fly binning to the Flow-Registration engine which are then concurrently compensated.

The average displacement of the last frames is used to initialize the lowest pyramid level of the displacement estimations in the following batch.

Reference frames can either be supplied directly or are computed from a specified set of frames, where the set of frames is pre-aligned with respect to the temporal average and then temporally averaged again. For this pre-alignment, the parameters *α* and *σ* are increased in size to make the result more robust under noise and reduce overfitting. The reference and data are normalized with respect to the 3D Gaussian filtered reference frames. Joint normalization is performed by default but channel-wise normalization is supported as well.

The displacements of each frame with respect to the reference on the lowpass filtered sequences are computed and the raw frames are then registered via backwards warping with bicubic interpolation, where out-of-bounds values are replaced with the values from the reference. To reduce quantization due to the interpolation, the results are stored with double precision as default but can also be stored with the precision of the input file.

To increase computational speed, the finest pyramid level for the OF calculation can be reduced in the options and the resulting low-resolution displacement field will be upsampled to the input image resolution in the final computation step. The Flow-Registration plugin implements this in the *Registration quality* setting, where only the highest quality setting calculates the solution on all levels. With settings *η* = 0.8 and minimum level of 6, we get almost tenfold faster computation time on the injection sequence with almost the same accuracy (PSNR −53.90 vs. −53.91). With those parameters, the highest resolution at which the displacements are estimated are given by 512 · 0.8^6^ = 134. Batch processing of multiple input files is possible via a batch processor class where a list of filenames is supplied that will be compensated either with individual references or the same reference.

To run the ImageJ / FIJI plugin, it needs to be installed via Plugins→Install Plugin which adds a Flow-Registration entry under Plugins. In its current version, the ImageJ / FIJI plugin can only be used for the compensation of shorter sequences depending on host system memory, the extension for virtual stacks is planned for the future. The MATLAB toolbox does not have such restrictions and can be used for the batch compensation of big data. For the MATLAB toolbox, a C++ compiler is required and the code has been tested for MATLAB R2018a onwards. The folder *demos* in the MATLAB toolbox contains scripts to reproduce the video results and quantitative results presented here as well as examples on how to use the code for different application scenarios. The *jupiter* demo compensates an amateur jupiter recording and demonstrates different aspects of multi-channel tiff and ROI processing as well as a potential application beyond the scope of neuroimaging.

## 3 Results

In this section, we first describe the benchmark dataset as well as the metrics we use for quantitative registration performance assessment. We then present the performance of our method in terms of registration quality and computational speed.

### 3.1 Benchmark datasets and metrics for evaluation

We used multiple datasets that have been recorded with the described setup previously. The dataset names indicate the imaging depth (see Figure 2 and Figure 3), where deeper layers (e.g. 5) correspond to lower SNR. For the layer 1-5 recordings, timepoints during movement onset which where contaminated with movement artifacts were selected from continuous recordings. For the datasets during drug and saline injections, we selected the time points around the injection events (e.g. compare Figure 1, (E)). From each dataset, we selected 2500 frames (80.9 s). A common approach for the evaluation of 30 Hz 2-photon microscopy data is temporal binning to increase the SNR. Therefore, our final benchmark dataset contains the data with binning over five frames (6.2 Hz, 500 frames total), as well as on a subset of the raw recordings, indicated by the suffix 30 Hz. For the dataset *saline injection*, in total 5 frames from the beginning and end of the experiment were excluded due to artifacts.

**Figure 2:**
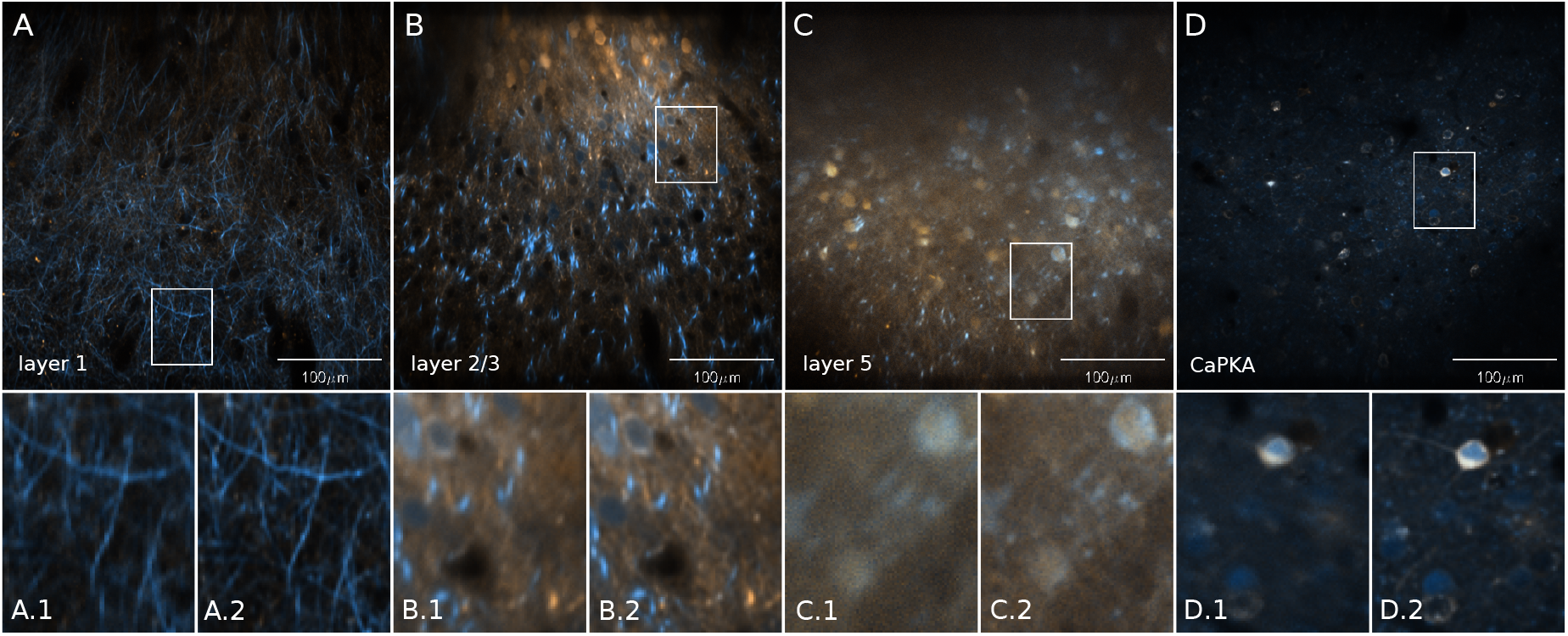
The benchmark datasets from layer 1 (A), layer 2/3 (B), layer 5 (C) and the CaPKA (D) dataset. Temporal average of raw recordings (A.1-D.1) and after application of Flow-Registration (A.2-D.2), both during the motion contaminated frames.

**Figure 3:**
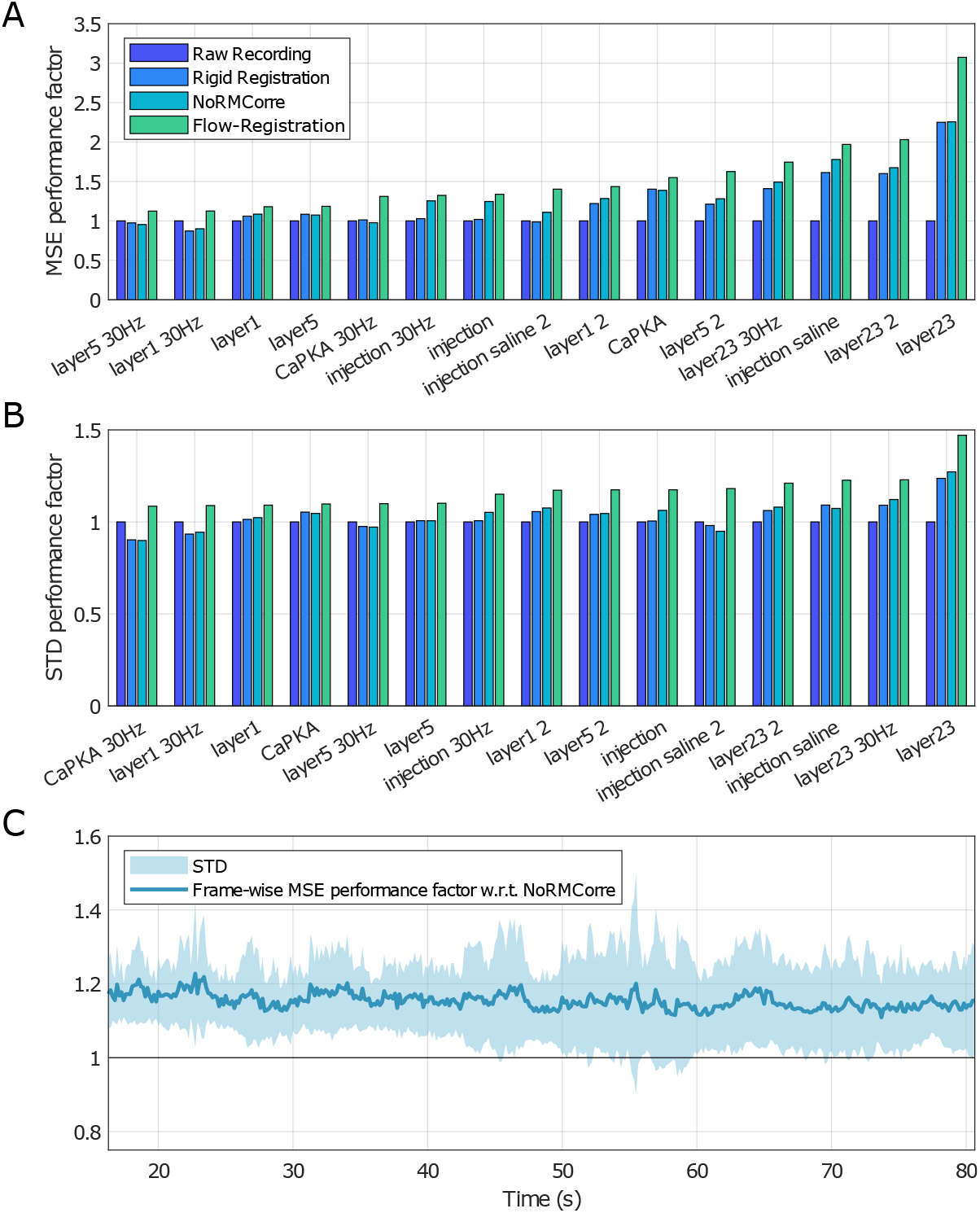
Application of Flow-Registration to 15 different datasets. We use 2-Photon recordings from layer 5, layer 2/3, layer 1 and three sequences during in vivo drug or saline injection at 6.2 Hz and 30.9 Hz. In all applied metrics, Flow-Registration performs consistently better than rigid registration and NoRMCorre. The metrics are averaged MSE (A) and STD (B) performance factors (see section 3.1) with respect to the raw recording. (C) displays the frame-wise MSE performance factor of Flowregistration with respect to NoRMCorre over all 6.2 Hz datasets with 500 frames (layer1 - layer 5, injection, injection saline 2; values > 1 indicate better performance of Flow-Registration). The frame-wise performance is significantly better for Flow-Registration (p < 0.00001, paired, two-sided Wilcoxon signed rank test).

**Figure 4:**
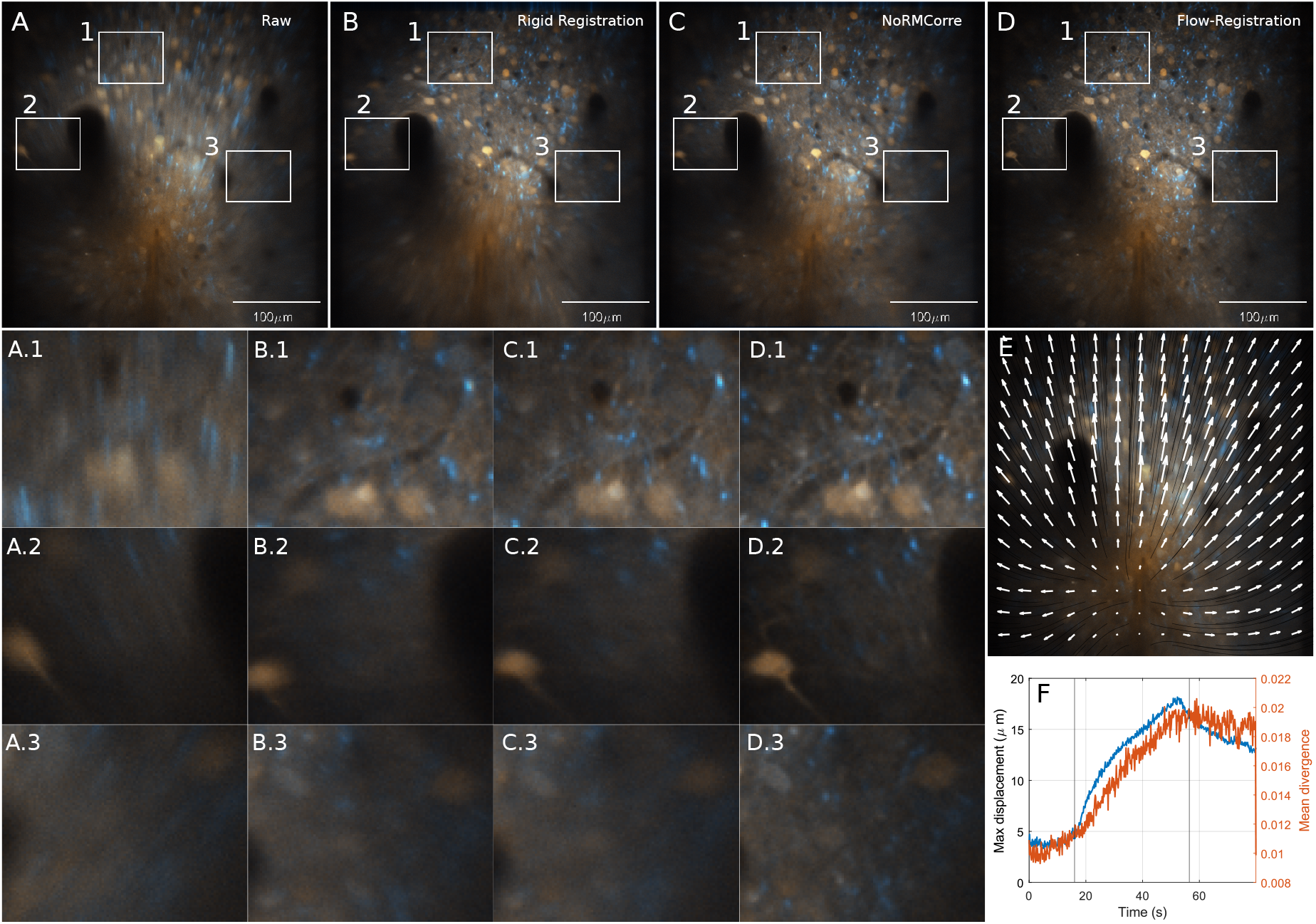
Qualitative results of the first in vivo saline injection sequence (dataset *injection saline*). Average of (A) raw images, (B) after rigid registration, (C) after registration with NoRMCorre, and (D) after Flow-Registration. The tissue expands from the injection point (E) resulting in large displacements as well as in a high divergence (F) in the displacement field. Flow-Registration can recover fine structures with much more detail, allowing region-of-interest selection along dendritic structures (blue), while the blur in (A-C) indicates residual motion.

**Figure 5:**
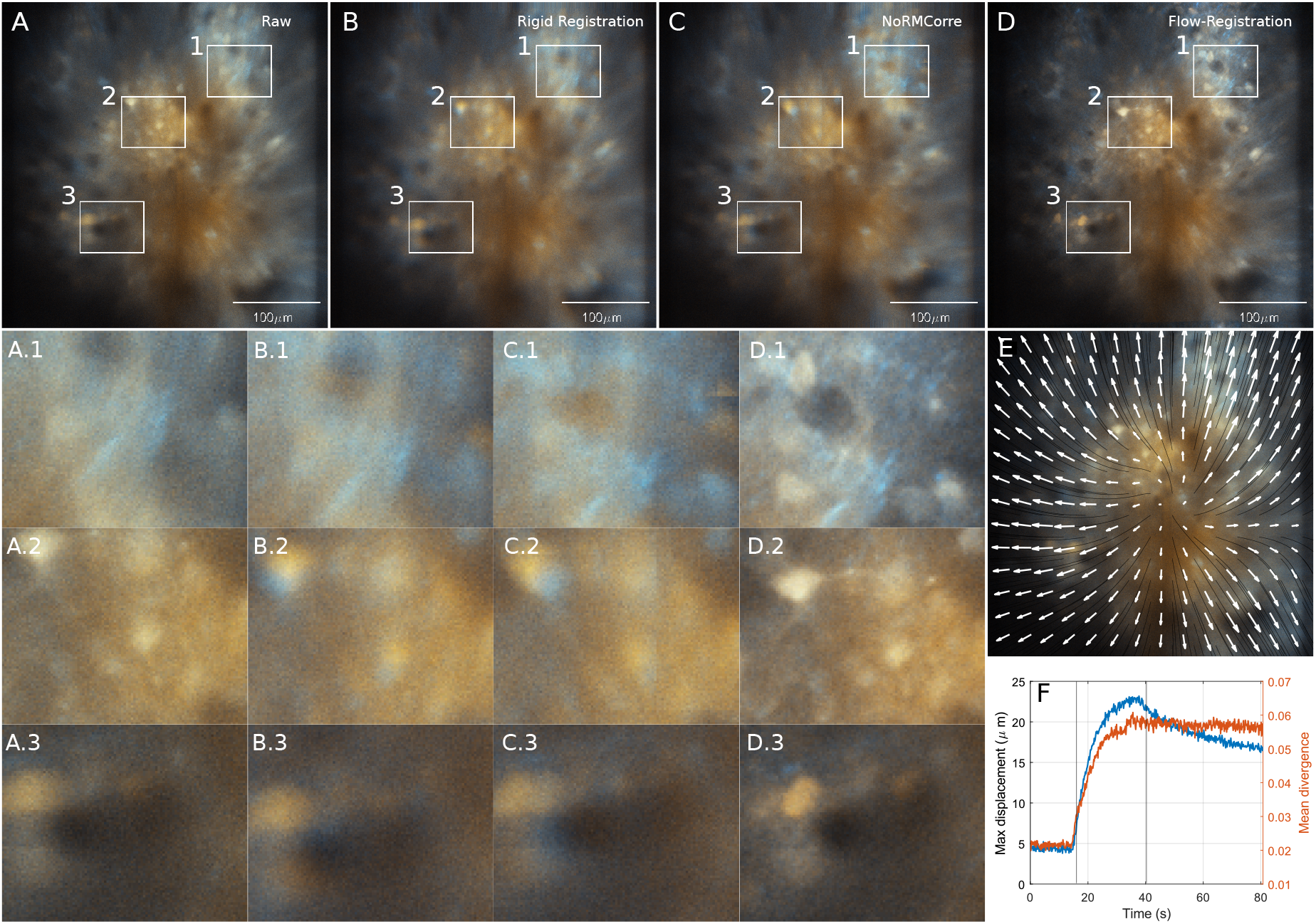
Qualitative results of the second in vivo saline injection sequence (dataset *injection saline 2*). Average of (A) raw images, (B) after rigid registration, (C) after registration with NoRMCorre, and (D) after Flow-Registration. The tissue expands from the injection point (E) resulting in large displacements as well as in a high divergence (F) in the displacement field. Flow-Registration can recover fine structures with much more detail, allowing region-of-interest selection along dendritic structures (blue), while the blur in (A-C) indicates residual motion.

Those data are real-world, low SNR datasets without ground truth displacements, therefore, metrics such as AEE or AAE are not applicable. The same holds true for perception based metrics due to the overall small movements and high image noise in the data. For the evaluation, we calculate reference based PSNR (see Table 1) and performance factors based on MSE as well as temporal STD (see Figure 3). To calculate the metrics, we first applied 2D Gaussian convolution with (*σ* = (3, 3)^⊤^). As reference, we used the temporal average over the reference frames from the motion compensation. The MSE and STD performance factors are then calculated as the fold change of the average MSE and STD values *MSE*(raw) · *MSE*(compensated)^-1^ and *STD*(raw) · *STD* (compensated)^-1^. They indicate the factor by which the MSE or STD value is higher on the raw sequence than the MSE or STD of each method (see Figure 3 (A) and (B)) or the factor by which the MSE value is higher for NoRMCorre when compared to Flow-Registration on a frame-wise basis (see Figure 3 (C)). We excluded all frames that contributed to the reference, as well as a boundary of 25 pixels. This boundary ensures that only valid regions are considered in the evaluation even with the large displacements of the injection sequences. Lowpass filtering reduced the influence of image noise on the results. For the compensation of all sequences, the first 100 frames (6.2 Hz) and 500 frames (30.9 Hz) were used as reference and supplied to the respective method.

**Table 1:** Average PSNR values on each dataset for the different methods. Flow-Registration consistently outperforms the other methods. PSNR has been calculated with respect to the maximum value 2^16^ and the images have been low-pass filtered before (see section 3.1) as well as SNR of the raw data (e.g. 30.9 Hz vs 6.2 Hz) have an impact on the PSNR differences between the datasets. The highest PSNR value in each row is put in bold.

|  | Raw | Rigid | NoRMCorre | Flow-Registration |
| --- | --- | --- | --- | --- |
| CaPKA | -10.10 | -8.81 | -8.82 | **-8.47** |
| CaPKA 30Hz | -14.22 | -14.59 | -14.64 | **-13.32** |
| injection | -57.35 | -56.73 | -54.95 | **-53.90** |
| injection 30Hz | -57.93 | -57.42 | -56.11 | **-55.35** |
| injection saline | -18.11 | -16.06 | -15.53 | **-15.01** |
| injection saline 2 | -17.70 | -17.69 | -17.11 | **-16.15** |
| layer1 | -4.54 | -4.37 | -4.28 | **-3.93** |
| layer1 2 | -8.74 | -8.19 | -8.01 | **-7.61** |
| layer1 30Hz | -11.17 | -11.77 | -11.65 | **-10.67** |
| layer23 | -19.19 | -16.44 | -16.40 | **-15.01** |
| layer23 2 | -7.01 | -5.35 | -5.11 | **-4.27** |
| layer23 30Hz | -23.08 | -21.78 | -21.49 | **-20.75** |
| layer5 | -6.14 | -5.83 | -5.87 | **-5.44** |
| layer5 2 | -7.54 | -6.95 | -6.76 | **-5.74** |
| layer5 30Hz | -12.55 | -12.66 | -12.75 | **-12.05** |

### 3.2 Registration performance

Flow-Registration consistently outperforms NoRMCorre [5] and rigid image registration on the benchmark datasets in terms of the reduced temporal STD and MSE ratios described in the previous section as well as *peak signal to noise ratio* (PSNR), see Figure 3 and Table 1. Frame-wise comparison of frames registered with Flow-Registration and NoRMCorre shows a significantly higher (p < 0.00001, paired, two-sided Wilcoxon signed rank test) performance of Flow-Registration, see Figure 3 (C). The graph indicates the factor, by which the NoRMCorre MSE is higher than the Flow-Registration MSE (i.e. > 1 indicates better performance of our method). Qualitatively, our method produces much crisper average frames compared to other methods during challenging motion events such as local drug injections with large intensity changes in the different channels, see Figure 1 and supplementary figures.

For a fast approximation of the solution, the minimum warping depth can be used to define the maximum resolution at which the solution is computed. We found around 10x speedup to be possible without reducing compensation quality (avg PSNR −53.90 with the precise and −53.91 with the approximated solution, min level = 6, compare supplemental Video 1).

In terms of computation time, the MATLAB toolbox is similar to existing methods and even faster with the approximated solution. On a single channel version of the 500 frame injection sequence (consumer workstation, 12 cores @ 3.8 GHz, 64GB of memory), our method (50 iterations, no pre-processing / IO, min level 0) takes 115 s and only 15 s with min level 6, while NoRMCorre (grid size 32, one iteration) takes 160 s and with grid size 16 even 763 s (setting used for the benchmark solution).

Flow-Registration was run for all datasets with default parameters and *α* = 1.5, for the 6.2 Hz datasets with *σ* = (1, 1, 0.1). and on the 30.9 Hz datasets with *σ* = (1, 1, 0.5)^⊤^. For the injection sequence, additionally the channel weight was set to (1.15, 0.85).^⊤^ The batch size was set to the size of the datasets.

## 4 Discussion

We have presented the new motion compensation method Flow-Registration and can report state-of-the-art registration results. The MATLAB implementation outperforms NoRMCorre in terms of registration quality as well as computation speed with comparable parameters. NoRMCorre is an established 2-photon neuroimaging motion compensation method and the default method in recent 2-Photon and calcium imaging suites such as CaImAn (2019) [22], EZcalcium (2020) [23] or Begonia (2021) [21] and can be considered the state of the art for the alignment of 2-photon videos. Our toolbox can produce output files that are compatible with CaImAn as well as with Begonia and can be easily extended for other frameworks and workflows via object oriented IO file readers and writers. Motion compensation methods often incorporate many regularization parameters that require difficult fine tuning for different motion statistics. For example, NoRMCorre has a total of 7 regularization and 2 subpixel refinement parameters [5]. Our method needs only a single regularization parameter and one parameter for presmoothing, while the results have native subpixel precision because of the continuous modeling.

Generally, a drawback of 2D motion compensation approaches for 2-photon imaging is the lack of z-shift correction. While there exist methods for high-speed, online 3D compensation [24], they require complicated setups and the current generation is limited to rigid 3D motion compensation — which might therefore benefit from refinement with a method such as Flow-Registration as well.

While our method does not aim for online processing, deep learning based approaches could unlock real time applications of our method. Even state-of-the-art self-supervised methods often require annotated data as initialization [25]. Flow-Registration has already been applied to many state-of-the-art 2-photon imaging recordings. The explicit, high-accuracy estimation of displacements with our method can be used for the generation of datasets for the training of efficient, deep learning based motion compensation methods in the future.

## 5 Conclusion

The solutions presented in this paper solve the pre-processing problem of motion contamination in 2-photon microscopy and multichannel video recordings with difficult, non-linear motion. Our core software design paradigm was the easy yet versatile integration into different workflows and toolboxes for 2-Photon imaging. We have developed a MATLAB toolbox that supports common file formats such as MDF, tiff image stacks, MATLAB mat files or HDF5 files in single file or multichannel configurations. The image IO is designed in a modular, object oriented way, such that the toolbox can easily be extended with new data formats and embedded pre-processing. The code is memory efficient and allows to compensate bigdata recordings with specified pre-processing methods and on-the-fly binning. The computationally heavy code is written in C++ which allows the implementation of Python wrappers in the future. The ImageJ / FIJI plugin builds on Imglib2 [26] library which supports most common image formats through the bio-formats plugin. The implementations support weighted, multichannel input with weight masks to enforce higher weight on the dataterm inside of ROIs.

The ImageJ / FIJI plugin is integrated with the MATLAB toolbox, such that the plugin can export parameters and reference frame configurations, that can be loaded as MATLAB bulk registration jobs.

Documentation, the MATLAB code for Flow-Registration and the ImageJ Plugin can be found on the GitHub repository https://github.com/phflot/flow_registration. The version used for the evaluations in this paper is supplied as supplemental code. It contains MATLAB scripts to reproduce the results reported here as well as the precompiled ImageJ / FIJI plugin.

## Acknowledgments

This work was partially funded by the Japan Society for the Promotion of Science (JSPS) with the Summer Program 2019 under grand number SP19305 and partially conducted at the *Optical Neuroimaging Unit* under Bernd Kuhn at the Okinawa Institute of Science and Technology Graduate University. The authors would like to thank Miles J. Desforges, Mohamed Eltabbal and Lars Haab for code testing and valuable feedback and discussions.

## Author contributions

**Philipp Flotho**: Software, Methodology, Conceptualization, Data Curation, Formal Analysis, Visualization, Writing-original draft; **Shinobu Nomura**: Data Curation, Investigation, Writing-review & editing, Validation; **Bernd Kuhn**: Conceptualization, Writing-review & editing, Supervision, Resources; **Daniel J. Strauss**: Conceptualization, Writing-review & editing, Supervision

## Data Availability Statement

The complete benchmark dataset used in this work is available as *2-Photon Movies with Motion Artifacts* on Dryad.

## A Parameter Selection and Data Presentation

### A.1 Parameter Selection

For Flow-Registration, parameters were chosen based on visual inspection. For NoRM- Corre, parameters from the NoRMCorre repository demo scripts where modified based on the statistics such as maximum displacements that have been calculated from the OF results. NoRMCorre incorporates different regularization approaches that makes it more robust under noise and implicitly define the properties of the compensated displacements. There are a total of 7 parameters for regularization and 2 for subpixel refinement against the single regularization parameter α and additional solver specific parameters for Flow-Registration, while subpixel accuracy is natively supported. We found it necessary to modify 4 of the NoRMCorre parameters for our datasets (see Table 2). Dynamic update of the reference was turned off due to the reference based metrics that were used for evaluation. In practice, on the injection sequences, this reduced drift, but slightly increased jitter at the same time.

**Table 2:**
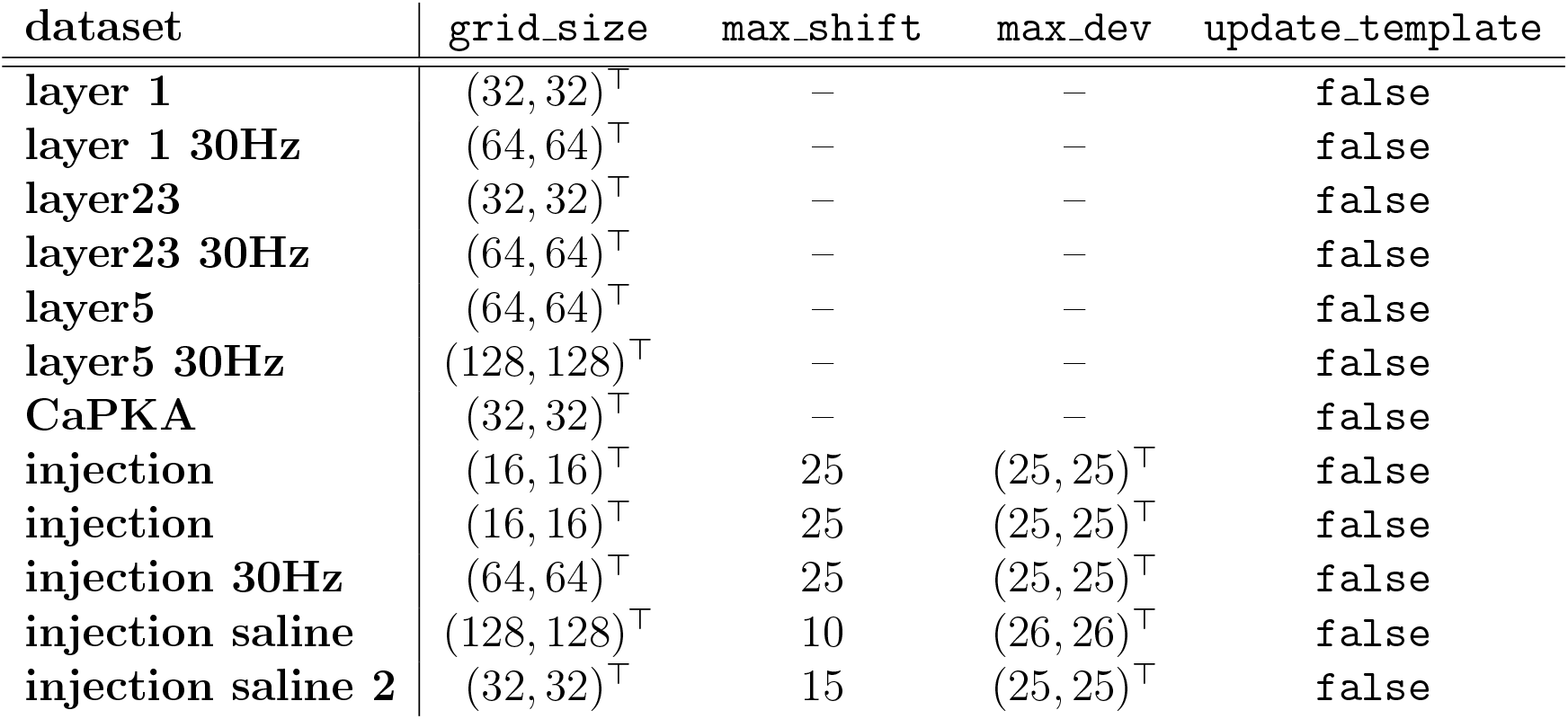
NoRMCorre parameters used for the compensation of the benchmark data. All unmentioned parameters have been kept in the default setting of the MATLAB code (October 2021). If the second version of the dataset is not mentioned, parameters match with the first version.

### A.2 False Color Representation

The microscopy images presented in this paper aim to visualize artifacts in the average frames caused by residual motion in two imaging channels. Therefore, we make use of an inclusive (linear) false color representation that allows distinction of the channels both with deuteranopia and protanopia. Given the minimum intensity values *l* = (*l*_1_, *l*_2_)^⊤^ and maximum intensity values *h* = (*h*_1_, *h*_2_)^⊤^ from the temporal average of the raw recordings, the color (*R,G,B*)^⊤^ at a given image position *f*(*x,y*) = (*p*_1_, *p*_2_)^2^ is calculated as

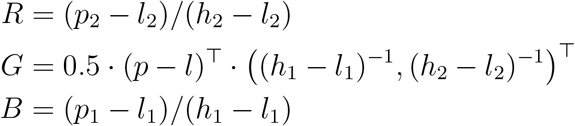

This means that the reference normalized red imaging channel is mapped onto *B*, the green channel onto *R* and *G* is the average of the other two. Therefore, turquoise areas correspond channel 2 of the input, orange areas to channel 1 and white areas indicate that both channels are active.

## B Supplemental Figures

## References

[1] Helmchen, F. & Denk, W. Deep tissue two-photon microscopy. Nat Methods 2, 932 (2005).

[2] Theer, P., Kuhn, B., Keusters, D. & Denk, W. Two-photon microscopy and imaging (2006).

[3] Horn, B. K. P. & Schunck, B. G. Determining optical flow. Artif Intell 17, 185–203 (1981).

[4] Greenberg, D. S. & Kerr, J. N. D. Automated correction of fast motion artifacts for two-photon imaging of awake animals. J Neurosci Methods 176, 1–15 (2009).

[5] Pnevmatikakis, E. A. & Giovannucci, A. Normcorre: An online algorithm for piecewise rigid motion correction of calcium imaging data. J Neurosci Methods 291, 83–94 (2017).

[6] Chen, Z., Jin, H., Lin, Z., Cohen, S. & Wu, Y. Large displacement optical flow from nearest neighbor fields. In Proc IEEE Comput Soc Conf Comput Vis Pattern Recognit, 2443–2450 (2013).

[7] Baker, S. et al. A database and evaluation methodology for optical flow. Int J Comput Vis 92, 1–31 (2011).

[8] Sun, D., Roth, S. & Black, M. J. Secrets of optical flow estimation and their principles. In Proc IEEE Comput Soc Conf Comput Vis Pattern Recognit, 2432–2439 (IEEE, 2010).

[9] Brox, T., Bruhn, A., Papenberg, N. & Weickert, J. High accuracy optical flow estimation based on a theory for warping. In Comput Vis ECCV, 25–36 (Springer, 2004).

[10] Bonnot, A. et al. Single-fluorophore biosensors based on conformation-sensitive gfp variants. FASEB J 28, 1375–1385 (2014).

[11] Nomura, S. et al. Combined optogenetic approaches reveal quantitative dynamics of endogenous noradrenergic transmission in the brain. iScience 23, 101710 (2020).

[12] Roome, C. J. & Kuhn, B. Chronic cranial window with access port for repeated cellular manipulations, drug application, and electrophysiology. Front Cell Neurosci 8, 379 (2014).

[13] Roome, C. J. & Kuhn, B. Voltage imaging with annine dyes and two-photon microscopy. In Multiphoton Microscopy, 297–334 (Springer, 2019).

[14] Zimmer, H. et al. Complementary optic flow. In Energy Minimization Methods Comput Vis Pattern Recognit, 207–220 (Springer, 2009).

[15] Hermosillo, G., Chef d’Hotel, C. & Faugeras, O. Variational methods for multimodal image matching. Int J Comput Vis 50, 329–343 (2002).

[16] Flotho, P. et al. Fast variational alignment of non-flat 1d displacements for applications in neuroimaging. J Neurosci Methods 353, 109076 (2021).

[17] Roome, C. J. & Kuhn, B. Dendritic coincidence detection in Purkinje neurons of awake mice. Elife 9, e59619 (2020).

[18] Bruhn, A. Variational optic flow computation: accurate modelling and efficient numerics. Ph.D. thesis, Saarland University (2006).

[19] Papenberg, N., Bruhn, A., Brox, T., Didas, S. & Weickert, J. Highly accurate optic flow computation with theoretically justified warping. Int J Comput Vis 67, 141–158 (2006).

[20] Bruhn, A., Weickert, J. & Schnörr, C. Lucas/Kanade meets Horn/Schunck: Combining local and global optic flow methods. Int J Comput Vis 61, 211–231 (2005).

[21] Bjørnstad, D. M. et al. Begonia—a two-photon imaging analysis pipeline for astrocytic ca2+ signals. Front Cell Neurosci 15, 176 (2021).

[22] Giovannucci, A. et al. Caiman an open source tool for scalable calcium imaging data analysis. Elife 8, e38173 (2019).

[23] Cantu, D. A. et al. Ezcalcium: open-source toolbox for analysis of calcium imaging data. Front Neural Circuits 14, 25 (2020).

[24] Griffiths, V. A. et al. Real-time 3d movement correction for two-photon imaging in behaving animals. Nat Methods 17, 741–748 (2020).

[25] Liu, P., Lyu, M., King, I. & Xu, J. Selflow: Self-supervised learning of optical flow. In Proc IEEE Comput Soc Conf Comput Vis Pattern Recognit, 4571–4580 (2019).

[26] Pietzsch, T., Preibisch, S., Tomančák, P. & Saalfeld, S. Imglib2 - generic image processing in java. Bioinformatics 28, 3009–3011 (2012).

